# Cryptic diversity and multiple origins of the widespread mayfly species group *Baetis rhodani* (Ephemeroptera: Baetidae) on northwestern Mediterranean islands

**DOI:** 10.1101/056556

**Authors:** Roberta Bisconti, Daniele Canestrelli, Roberta Tenchini, Carlo Belfiore, Andrea Buffagni, Giuseppe Nascetti

**Affiliations:** Dipartimento di Scienze Ecologiche e Biologiche, Università della Tuscia. Viale dell’Università s.n.c., I-01100 Viterbo, Italy.; CNR – IRSA, Water Research Institute, Via del Mulino 19, 20861 Brugherio (MB), Italy

**Keywords:** Island biogeography, Mediterranean basin, *Baetis*, Tyrrhenian islands, Vicariance, Dispersal

## Abstract

How the often highly endemic biodiversity of islands originated has been debated for decades and it remains a fervid research ground. Here, using mitochondrial and nuclear gene sequence analyses, we investigate the diversity, phylogenetic relationships, and evolutionary history of the mayfly *Baetis gr. rhodani* on the three largest north-western Mediterranean islands (Sardinia, Corsica, Elba). We identify three distinct, largely co-distributed, and deeply differentiated lineages, with divergences tentatively dated back to the Eocene-Oligocene transition. Bayesian population structure analyses reveal a lack of gene exchange between them, even at sites where they are syntopic, indicating that these lineages belong to three putative species. Their phylogenetic relationships with continental relatives, together with the dating estimates, support a role for three processes contributing to this diversity: (1) vicariance, primed by microplate disjunction and oceanic transgression; (2) dispersal from the continent; and (3) speciation within the island group. Thus, our results do not point toward a prevailing role for any of the previously invoked processes. Rather, they suggest that a variety of processes equally contributed to shape the diverse and endemic biota of this group of islands.

## Introduction

Which processes best explain patterns of diversity on islands has been a central topic of island biogeography since its inception, and it continues to be a hot-topic for research and debate (Whittaker and Fernàndez-Palacios, 2007). The geological origin of an island system has been traditionally regarded as a major factor in explaining the structure of its biota. Accordingly, the highly distinctive biota of continental islands was often interpreted as the long-term product of divergence in isolation following habitat fragmentation, more or less complemented by neoendemic forms (Watson, 2009). This traditional view has been substantially enriched thanks to the advent of sharper research tools (e.g. population genetic, molecular phylogenetic, and phylogeographic approaches). Several of the best-studied island systems of continental origin have appeared to have an ‘oceanic’ nature, that is, they are dominated by neoendemic forms (e.g. Grandcolas et al., 2008; Goldberg et al., 2008). Vicariance and long-distance dispersal, long regarded as opposing explanations (Heads, 2009), are now increasingly perceived as often interacting processes in the assembly of an insular biota (Lomolino et al., 2010). In addition, microevolutionary processes priming divergence among populations within island have been recognized as substantial contributors to the biodiversity of islands, regardless of those islands’ geological origin and the mode of species settlement on them (e.g. Bisconti et al., 2013a, 2013b; Emerson et al., 2006; Holland and Hadfield, 2002; Thorpe and Malhotra, 1998; Villacorta et al., 2008; Wallis and Trewick, 2009). Although a few iconic island systems have dominated the scene of such reappraisal, it is noteworthy that as more islands and taxa are investigated, the range of processes seen as contributing to the assembly and diversity of current insular biota becomes wider (e.g. Bauzà-Ribot et al., 2011; Bisconti et al., 2011a; Salvi et al., 2014).

The Tyrrhenian (northwestern Mediterranean) islands belong to the largest continental fragment within the Mediterranean basin. This fragment originated by detachment from the Iberian Peninsula, the counterclockwise rotation of the microplate (initiated 21 to 15–18 Ma ago; Gattaccecca et al., 2007), and subsequent oceanic spreading (Brandano & Policicchio, 2012), while the disjunction of the two main islands, Sardinia and Corsica, was completed about 9 Ma ago (Alvarez, 1972; Bellon et al., 1977; Bonin et al., 1979). Subsequent land connections between islands and with the continent formed repeatedly, as a consequence of Mediterranean sea-level oscillations (Hsü et al., 1973; Krijgsman et al., 1999; Shackleton et al., 1984; Van Andel and Shackleton, 1982). This island system shows an extremely wide range of climates, landscapes, and habitats, and both its paleoclimates and its palaeoenvironments have been investigated in detail (Vogiatzakis et al., 2008). Consequently, in light of the wealth of background information available, it offers an ideal system for exploring both traditional and new hypotheses about the processes leading to the assembly and uniqueness of continental island biotas, an opportunity that has not escaped attention in the recent past.

Several hypotheses on the origin of the Tyrrhenian islands biota have been put forward during the last century. At first, it was proposed that such a distinctive biota formed via hologenesis, a process implying a prominent role for vicariance events (Monti, 1915; Monterosso, 1935; Luzzatto et al., 2000). Subsequently, more emphasis was given to dispersal, and a three-step process of colonization was hypothesized (Baccetti, 1964): a pre-Miocene step through a supposed land bridge with the Baetic region (southern Spain); a Miocene step which brought warm-temperate species from North Africa and Italy; and a Quaternary step which brought temperate species from the Apennines (Italy) via a land bridge connecting Tuscany with northern Corsica. Once the theory of plate tectonics spread (Alvarez, 1972), a reappraisal of vicariance as the leading force occurred, particularly to explain the most distinctive (endemic) components of this insular biota (Baccetti, 1980). With the advent of molecular phylogenetic approaches, data on several taxonomic groups began to accumulate (Grill et al., 2007; Ketmaier and Caccone, 2013), leading to a reconsideration of dispersal events, although vicariance events primed by the microplate disjunction are still regarded by several authors as the main triggers of endemic species formation on the islands (Ketmaier and Caccone, 2013). Finally, in more recent times, the application of the phylogeographic toolbox to the study of evolutionary histories within-islands (i.e. post-settlement), is shedding light on a plethora of previously unappreciated evolutionary processes contributing to the diversity of the island biota above and below the species level (Bisconti et al., 2011a, 2011b; Bisconti et al., 2013a, 2013b; Falchi et al., 2009; Gentile et al., 2010; Ketmaier et al., 2010; Salvi et al., 2009; Salvi et al., 2010).

In this paper, we focus on mayflies of the *Baetis rhodani* (Pictet, 1843) species group, which is one of the most common mayflies in the Western Palaearctic region, and one of the most abundant insects in freshwater running environments (Brittain, 1982). Phylogenetic and phylogeographic investigations of this species group have proliferated in recent years. In spite of the homogeneity of morphological characters, such studies have revealed unexpectedly deep divergences. Williams and colleagues (2006) investigated the genetic structure of *B. rhodani* populations in the Western Palaearctic, and identified multiple divergent lineages, probably belonging to distinct cryptic species. Lucentini et al. (2011), expanded the analysis to other European countries (UK and Italy), increasing the number of lineages. More recently, populations from the Canary Islands have also been investigated (Rutschmann et al., 2014), again showing deep divergences and several endemic lineages, likely resulting from a complex evolutionary history. Unfortunately, these studies were based on a single mitochondrial DNA marker (mostly cytochrome oxidase I), preventing sound inferences about the species status and the evolutionary processes involved. Nonetheless, results of these studies point to these mayflies as potentially excellent models for evolutionary and historical biogeographic studies in the Western Palaearctic.

Within the Tyrrhenian islands, a single endemic species, *Baetis ingridae* (Thomas and Soldàn, 1987) has been described from the *B. rhodani* group, based on a sample from northeastern Corsica. This species shows subtle morphological differences with respect to continental *B. rhodani.* Their larvae are very similar, with differences affecting colour patterns, the shape of the femoral bristles, and the shape and number of the teeth on the inner margin of the paraproct (Bauernfeind and Soldàn, 2012). So far, the diversity (including genetic diversity) and phylogenetic relationships of this mayfly within the Mediterranean islands is still virtually unexplored.

Here we employ a set of nuclear (nDNA) and mitochondrial (mtDNA) gene regions with the aim of investigating (1) the diversity of the *B. rhodani* species group in the Tyrrhenian islands; (2) the phylogenetic relationships with its continental relatives; (3) its evolutionary history, with special reference to the mode of settlement on islands; and ultimately (4) to aid shedding more light on the historical origin of these islands’ highly endemic diversity.

## Materials and Methods

### Sampling

Larval individuals of the *Baetis rhodani* species group were collected from aquatic habitats at 28 localities: 23 localities from the two main Tyrrhenian islands (Sardinia and Corsica); 2 samples from the Elba island, which is close to the northeastern side of Corsica and shows faunal and floral affinities with that island; and 3 samples from peninsular Italy (i.e. the nearest mainland), added to this study for comparative purposes. Taxonomic assignment of the sampled individuals to the *B. rhodani* species groups was carried out using well established morphological characters of diagnostic value (Müller-Liebenau, 1969). The geographic location of the sampling sites and sample details are shown in Table 1 and Figure 1B. Individuals of the *Baetis vernus* species group (sampling coordinates: 42° 12.72’ N; 12°24.95’ E) were added to the sampling for use as an outgroup in phylogenetic analyses. The sampled individuals were transported to the laboratory and preserved in 95% ethanol until DNA extraction.

**Table 1.**
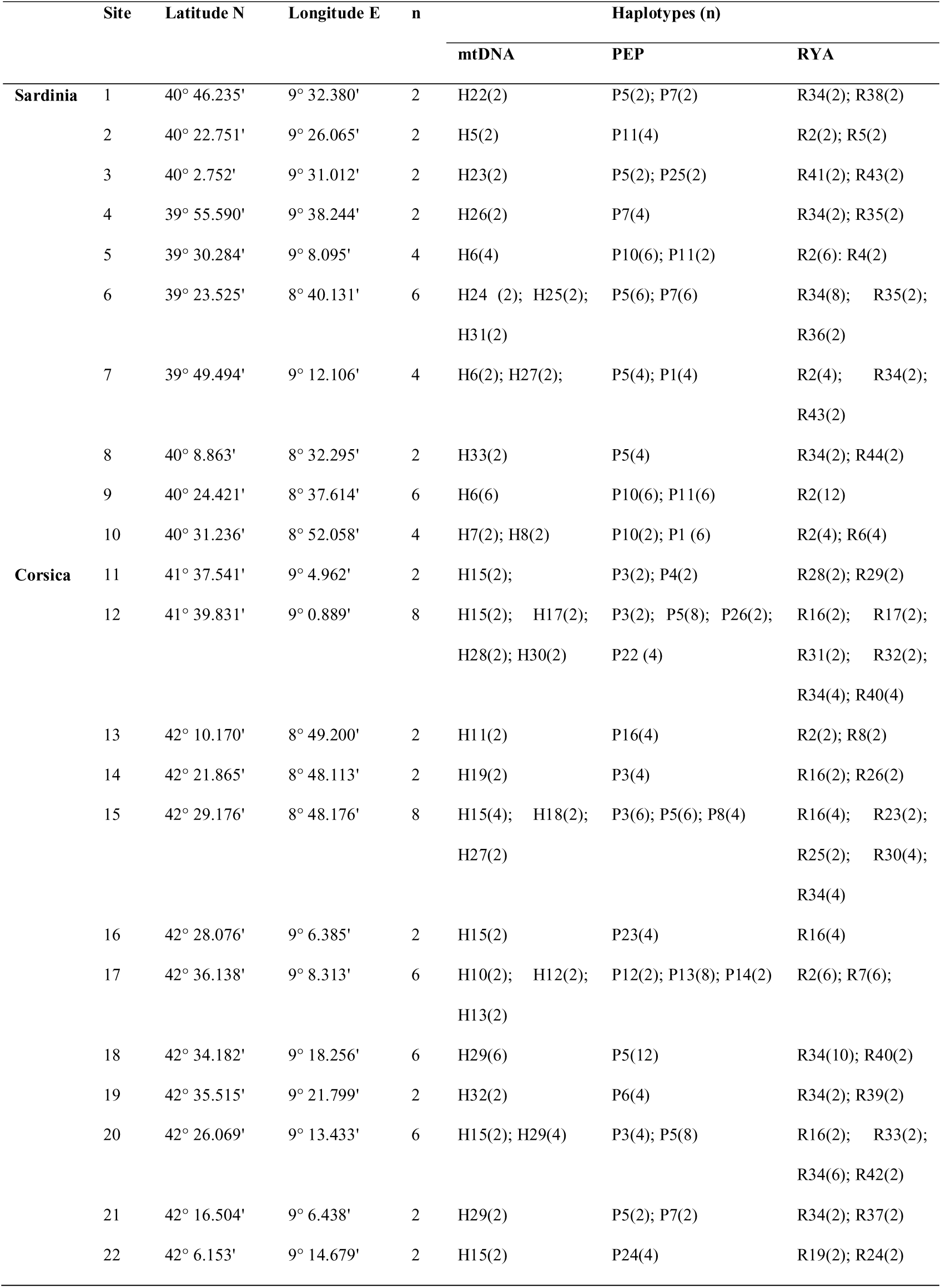
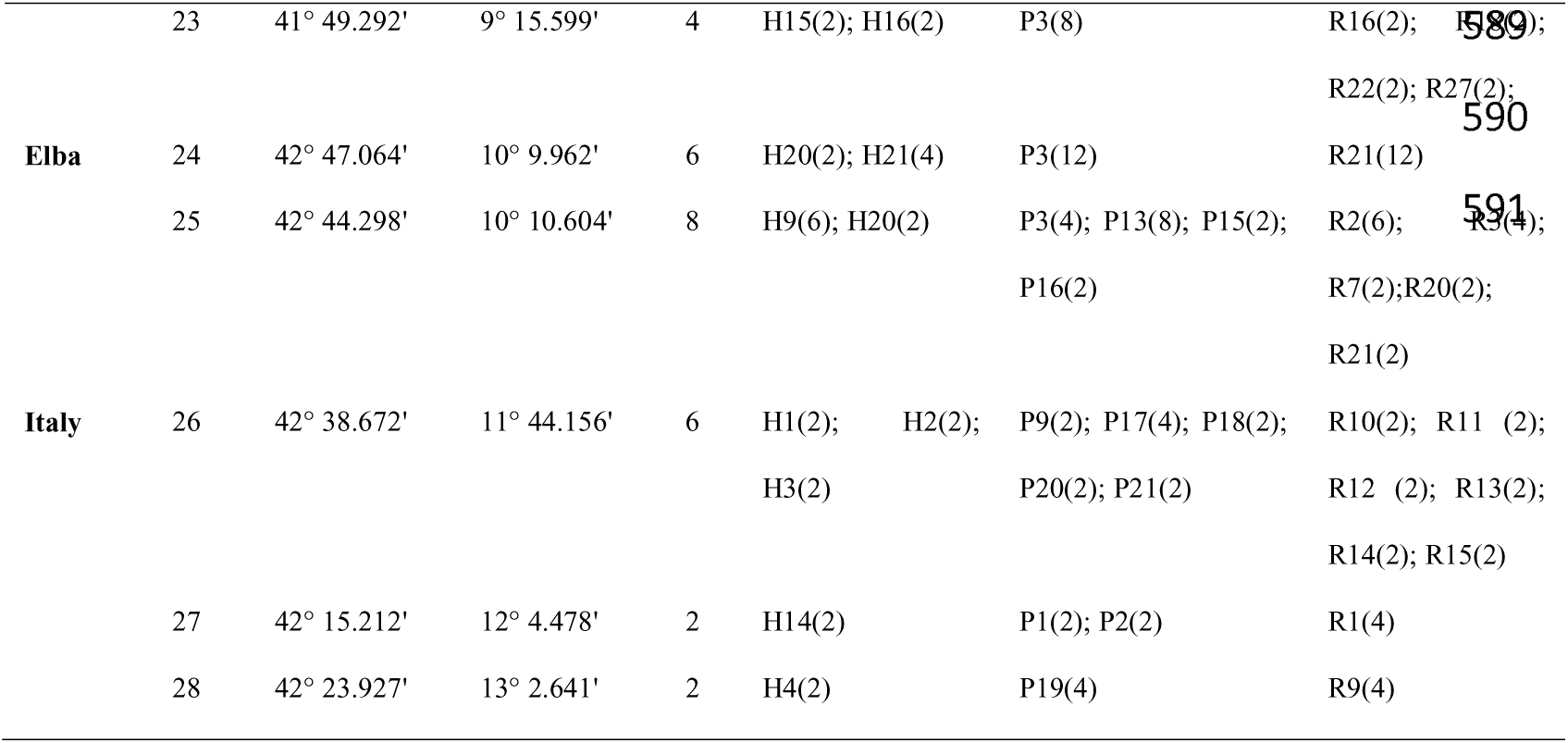
Geographical location and sample size of the 28 population samples of the *Baetis rhodani* species group used in this study. The geographic distribution of the haplotypes found and their abundance among populations is also given for each gene marker analysed.

**Figure 1.**
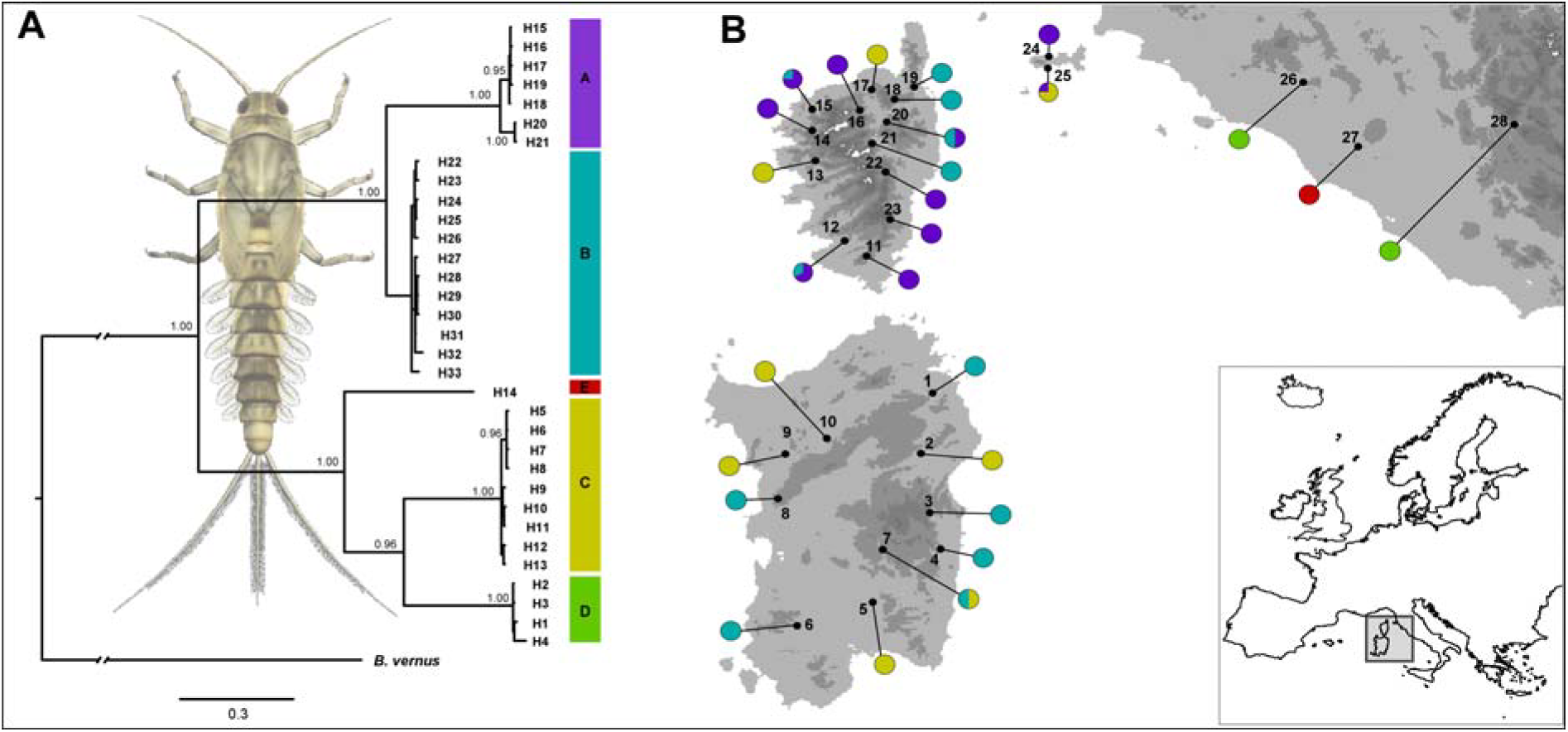
**(A)** Bayesian phylogenetic tree based on the 33 mtDNA haplotypes found in *Baetis rhodani* populations from the Tyrrhenian islands and neighbouring areas, estimated using MRBAYES. Posterior probabilities are shown at the nodes when ≥0.90. **(B)** Geographic locations of the 28 *Baetis rhodani* populations sampled, numbered as in Table 1. Pie-diagrams show the geographic location and frequency within populations of the main mtDNA haplogroups found.

### DNA extraction, amplification, and sequencing

We extracted whole genomic DNA using proteinase K digestion followed by a standard phenol-chloroform protocol with RNase treatment (Sambrook et al., 1989). Polymerase chain reactions (PCR) were carried out to amplify fragments of two mitochondrial genes (cytochrome oxidase 1, hereafter referred to as CO1; NADH dehydrogenase subunit 1, hereafter referred to as ND1), and two nuclear genes (phosphoenol pyruvate carboxykinase, hereafter referred to as PEPCK; ryanodine receptor 44f, hereafter referred to as RYA). PCR primers and protocols for the gene fragments CO1 and PEPCK were drawn from the literature (Simon et al. 1994; Pereira-da-Conceicoa et al., 2012). PCR primers for the ND1 gene fragment were designed using PRIMER 3, based on the complete genome alignment of *Parafronurus youi*, *Ephemera orientalis*, and *Baetis* sp., downloaded from the GenBank database (accession numbers EU349015.1, EU591678.1, and GU936204.1, respectively). For the RYA gene fragment, the PCR procedure was newly developed for the present study using the EvolMarkers bioinformatic tool (Li et al., 2010). With this tool, we searched for exon-primed intron-crossing (EPIC) markers, using annotated genomes of *Drosophila melanogaster* (‘query’), *Aedes aegypti* (‘subject’), and *Apis mellifera* (‘subject’), since annotated genomes of mayflies were not available. PCR primers were designed for 10 putative EPIC markers using PRIMER 3. After preliminary PCR trials, we focused on five putative markers which were consistently amplified in a sample of 10 randomly selected individuals, and which provided single and clean PCR bands in a standard (1.5x) agarose-gel electrophoresis. Among these markers, the RYA gene fragment returned high-quality sequence electropherograms, and was thus retained and used to analyse the entire pool of sampled individuals.

PCR cycling conditions were the same for all genes: 5 min at 92°C followed by 30 cycles of 1 min at 92°C, 1 min at an annealing temperature specific for each gene, and 90 s at 72°C, followed by a single final step of extension of 10 min at 72°C. Amplifications were conducted in 25 μL containing the following: MgCl_2_ (2.5 mM), the reaction buffer (1X; Promega), 4 dNTPs (0.2 mM each), 2 primers (0.2 μM each), the enzyme Taq polymerase (1 unit; Promega), and 2 μl of DNA template. PCR primers and annealing temperature specific for each gene fragment analysed are given in Table S1.

PCR products were purified and sequenced by Macrogen Inc. (http://www.macrogen.com) using an ABI PRISM 3700 sequencing system. The electropherograms were visually checked using CHROMAS 2.31 (Technelysium ltd.). All sequences were deposited in the GenBank database. Multiple sequence alignments were produced using CLUSTALX (Thompson et al., 1997), under the default settings.

### Data analysis

Sequence variation, substitution patterns, and divergence between haplotypes and between the main haplotype groups were assessed using MEGA 5.1 (Tamura et al., 2011).

Since this study is the first to investigate patterns of genetic diversity and affinities of mayflies of the *B. rhodani* group throughout the Tyrrhenian islands, we began our data exploration by verifying their purported endemicity, that is, whether they belong to well supported and divergent clades, geographically restricted to the Tyrrhenian islands. It is worth emphasizing, however, that we refrained from further interpreting the results of this analysis, since it relied on a final alignment of limited extension (see Results) and on a single gene fragment. This task was undertaken through a comparison of our CO1 data (the most widely used genetic marker in previous studies) with previously published sequences of *B. rhodani.* To this end, we retrieved one representative haplotype sequence of each main clade previously observed by Williams et al., (2006), Lucentini et al., (2011), and Murria et al., (2014), in order to have the widest possible coverage of continental Europe, and we built a Bayesian phylogenetic tree as described below.

Gene trees for the CO1 dataset, as well as for the concatenated mtDNA, PEPCK, and RYA datasets, were estimated by means of the Bayesian inference procedure implemented in MRBAYES v.3.2.1 (Ronquist et al., 2012). The nuclear gene sequences were phased using the software PHASE 2.1 (Stephens et al., 2003) with the default settings, while the possible occurrence of recombination was assessed using the pairwise homoplasy index (PHI statistic, Bruen et al., 2006) implemented in SPLITSTREE v.4.11 (Huson and Bryant, 2006). The best partitioning strategy for the concatenated mtDNA was selected by means of PARTITIONFINDER v1.0.1 (Lanfear et al., 2012), using the Bayesian information (BI) criterion and the ‘greedy’ search strategy. This analysis suggested TrN + I + G as the best model for this dataset. The best-fit model of nucleotide substitution for nuclear gene fragments (nDNA) was selected among 88 alternative models using the BI criterion implemented in JMODELTEST 2.1.3 (Darriba et al., 2012). This analysis indicated TrNef + G and K80 + G as the best-fit substitution models for the PEPCK and the RYA genes, respectively. For gene tree estimations, all the analyses consisted of two independent runs of four Markov chains, each run consisting of 10 million generations sampled every 1000 generations. Stationarity of the analyses was verified by examining log-likelihood scores using TRACER v1.6 (Rambaut et al., 2014), and chain convergence between simultaneous runs was evaluated by means of the average standard deviation of split frequencies (all values <0.001). For the nuclear datasets, which showed substantially lower levels of variation than the mtDNA dataset (see Results), gene trees were converted into haplotype networks using HAPLOTYPEVIEWER (Guidon et al., 2004).

Nuclear DNA data were further analysed using the non-hierarchical Bayesian clustering procedure implemented in BAPS 6.0 (Corander et al., 2008), which allows us to identify the best clustering option based on multilocus datasets, to assign individuals to clusters, and, importantly, to identify individuals of mixed ancestry among groups. We ran five replicates of this clustering analysis to check its consistency, based on the ‘clustering with linked loci’ option, the ‘independent loci’ model, and with the maximum number of clusters (KMAX) set to 20. The admixture analyses were run for 200 iterations, with 100 reference individuals and 50 iterations per reference individual.

Finally, we used the Bayesian inference procedure implemented in *BEAST (Heled and Drummond, 2010), in order to co-estimate individual gene trees for the four gene fragments analysed (CO1, ND1, RYA, PEPCK), and to embed them into a single multilocus species-tree. With this method, terminal taxa have to be indicated a priori, and so we used the results of previous phylogenetic inferences based on mtDNA and the non-hierarchical cluster analysis of nDNA data. Substitution, clock, and tree models were unlinked, and each dataset was assigned the appropriate ploidy level and the substitution model previously estimated by means of PARTITIONFINDER and JMODELTEST. The analysis was run using the Yule species-tree prior, and a strict molecular clock model was assumed. In order to get an approximate temporal context for the main divergences inferred, and in the absence of internal calibration points or fossil records, we used the CO1 specific substitution rate of 0.0115 substitutions per site per Ma (Brower, 1994) which has been extensively used to date divergences in insects, including mayflies, and which falls in the middle of the ranges estimated for this gene fragment in insects. Substitution rates for the remaining gene fragments were estimated using the CO1 specific rate as a reference, and the 1/x prior distribution. Five independent *BEAST analyses were performed to check for convergence, each run for 100 million generations, and sampling every 10,000. Convergence of the analyses, the appropriate number of samples to discard as burn-in (10%), stationarity, and the achievement of appropriate ESS values (≫200) were inspected using TRACER. The final species tree was estimated as a maximum clade credibility tree (MCC) summarizing results of the post-burn-in trees, with nodes annotated with ages and the corresponding 95% highest posterior densities (HPD), by means of the BEAST module TREEANNOTETOR. In addition, the entire posterior distribution of species-trees obtained with *BEAST was inspected using DENSITREE.

## Results

For the 112 individuals of the *B. rhodani* species group used in this study, we obtained fully resolved sequences (i.e., without missing data) for all the gene fragments analysed. Sequence length was as follows: 413 bp for CO1, 673 bp for ND1, 290 bp for PEPCK, and 178 bp for RYA. Within the combined mtDNA dataset (overall 1086 bp), 496 variable positions were found (422 excluding the outgroup), of which 390 were parsimony informative (381 excluding the outgroup). No indels, stop codons, nonsense codons, or heterozygous positions were observed within the CO1 and ND1 genes, consistent with the expectation for coding regions of mitochondrial origin. Within the nuclear gene sequences, indels were found of length 1-3 bp. For all downstream analyses, indels were removed from the alignments. The PEPCK gene presented 54 variable positions of which 45 were parsimony informative (41 and 32, respectively, excluding the outgroup), whereas the RYA gene showed 76 variable positions of which 61 were parsimony informative (54 and 32, respectively, excluding the outgroup). No statistically significant indications of recombination events were found with the PHI tests carried out on the nuclear gene fragments.

A CO1 alignment of length 412 bp was generated by pooling our CO1 sequence with those retrieved from the literature. The Bayesian tree estimation carried out on this dataset (Figure S1) clearly identified our individuals as belonging to 3 well supported clades (clades A, B, C) geographically restricted to the northwestern Mediterranean islands, and divergent from all other clades. Two of these clades (A, B) showed sister relationships, whereas the third clade (C) was nested within continental haplotypes. Thus, we found no evidence against the purported endemicity of the insular representatives of the *B. rhodani* species group, although they did not appear to be a monophyletic group.

The Bayesian phylogenetic analysis performed on the concatenated mtDNA (Figure 1A) identified a well resolved topology, with most nodes having a posterior probability ≥0.99. Five main clades were found: 3 observed within the Tyrrhenian islands (clades A, B, and C, as above), and 2 only found on the neighbouring mainland (clades D, E). Within clade A, two subclades were observed, one geographically restricted to Elba (haplotypes H20, H21), the other restricted to Corsica. Clade B was observed both in Corsican and Sardinian samples, whereas it was not observed on Elba. Instead, haplotypes of clade C were carried by individuals from all the sampled islands. Clades A, B, and C together were not monophyletic, the latter being nested within Italian clades. The mean uncorrected sequence divergence among clades (Table 2) ranged between 0.12 (clade A vs. B) and 0.269 (clade A vs. D), whereas mean within-clade divergence ranged between 0.006 (clade C) and 0.011 (clade A).

**Table 2.** Mean pairwise uncorrected sequence divergence among (below diagonal) and within (diagonal) the five main haplotype groups identified in the concatenated mtDNA dataset. Standard errors of the between-group divergence estimates are given above the diagonal. Haplotype groups are encoded as in Figure 1.

|  | **A** | **B** | **C** | **D** | **E** |
| --- | --- | --- | --- | --- | --- |
| **A** | **0.011** | 0.010 | 0.013 | 0.014 | 0.014 |
| **B** | 0.135 | **0.008** | 0.013 | 0.014 | 0.015 |
| **C** | 0.263 | 0.238 | **0.006** | 0.012 | 0.012 |
| **D** | 0.280 | 0.256 | 0.166 | **0.009** | 0.012 |
| **E** | 0.264 | 0.249 | 0.194 | 0.190 | **n/c** |

The non-hierarchical cluster analysis of the nDNA dataset carried out with Baps indicated K = 4 as the option best fitting the data (posterior probability = 1.00). Individuals from insular samples were assigned to three distinct clusters, fully corresponding to clades A, B, and C identified from the mtDNA dataset. Contrary to what was previously found from the mtDNA, all individuals from the mainland were grouped into a single cluster. Moreover, with this analysis, no individuals of mixed ancestry between clades were found, even within population samples where distinct mtDNA clades and nuclear clusters were observed to be syntopic (samples 7, 12, 20, 25).

The phylogenetic networks built among haplotypes found at the nuclear gene fragments PEPCK and RYA are shown in Figure 2. Within both networks, haplotypes from individuals carrying the mtDNA clades A, B, and C can be identified as groups of closely related nDNA haplotypes. Heterozygote individuals carrying haplotypes from two of these groups were not observed, not even within sites of syntopy. Haplotype from these groups were never observed In addition, a closer affinity between the nDNA group C (yellow in Figure 2) and the Italian haplotypes was observed. However, the genealogical relationships among Italian haplotypes and between them and haplotypes of group C appeared much less clearly resolved than with the mtDNA dataset. Within the insular populations where the three haplogroups from distinct groups groups were found, heterozygotes for haplotypes from the three distinct groups.

**Figure 2.**
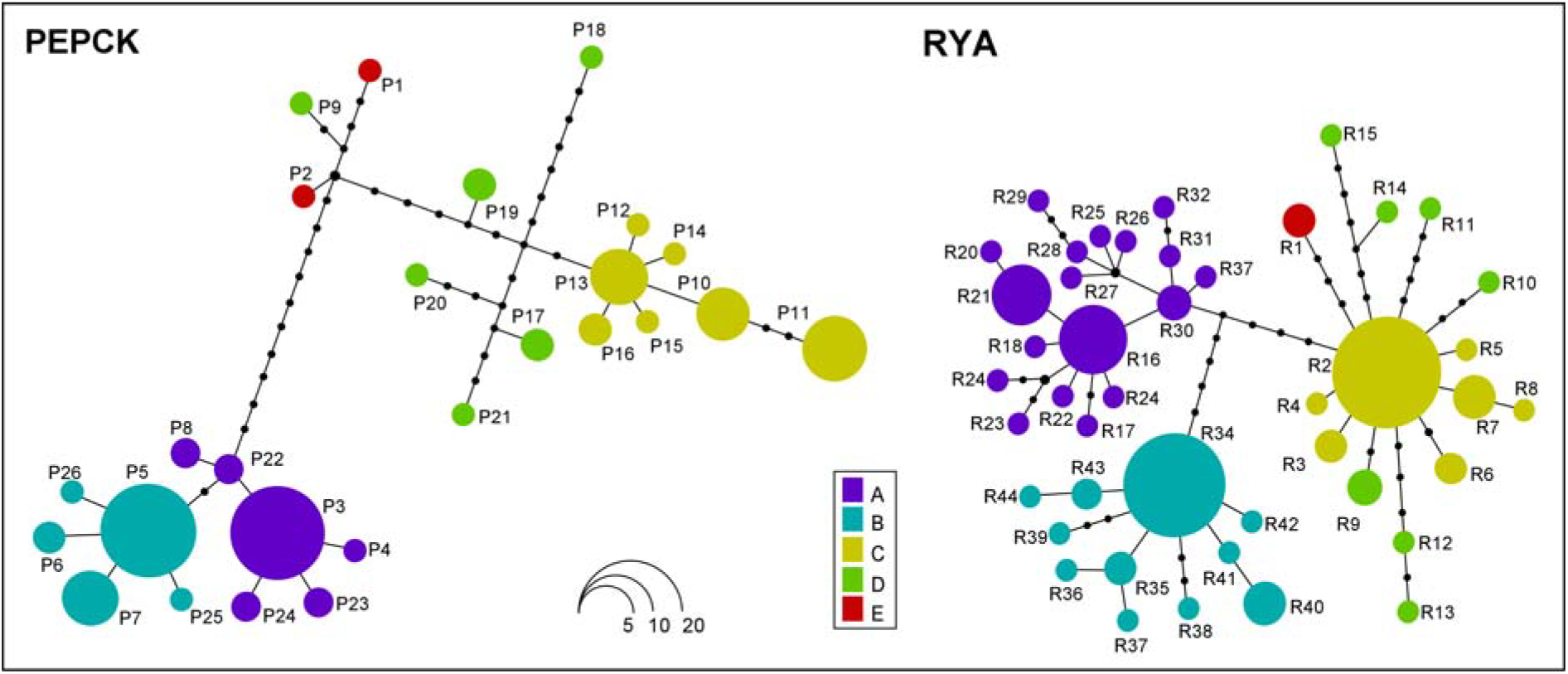
Phylogeographic networks generated with HAPLOTYPEVIEWER, based on the Bayesian phylogenetic trees estimated for each nDNA gene fragment using MRBAYES. Circle size is proportional to the frequency of the corresponding haplotype across the dataset. Each individual was given the same colour as the respective mtDNA haplogroup, for comparative purposes.

The multilocus species-tree analysis with *BEAST was carried out using three alternative strategies for the a priori definition of terminal taxa. First, following results based on the CO1 dataset, we added to our multilocus dataset two individuals from the closest relatives of our insular clades. We defined a set of 7 ingroup taxa, 5 resulting from the Bayesian analysis of the concatenated mtDNA data, plus 2 groups retrieved from the literature (G1-G2 [Lucentini et al., 2011] and G5-Brho2hap8 [Lucentini et al., 2011; Murria et al., 2014], here referred to as clades G and H). The *BEAST analysis was then run with the non-CO1 data from these 2 latter groups encoded as missing. Second, the same analysis was re-run after the exclusion of CO1 sequences derived from the literature (i.e. with 5 terminal taxa). Third, individuals were assigned to four terminal taxa defined according to results of the non-hierarchical cluster analysis of the nDNA data, that is, with the mainland samples grouped together. With 7 groups (see Figure 3), the *BEAST analysis confirmed with high support a closer affinity between clade C and the continental clade G (rather than between clade C and the other insular clades). The divergence between the two main clades was estimated to have occurred at 31.4 Ma (95% HPD: 21.0–42.4), whereas the divergence between the two insular clades A and B was dated at 7.4 (95% HPD: 4.6–10.3). Divergence between clade A + B and its closest continental relative (H) was estimated at 19.9 (95% HPD: 12.0–29.3). Instead, the divergence between the insular clade C and clade G was dated at 2.8 (95% HPD: 0.5–4.8). Carrying out the analysis with 4 terminal taxa did not appreciably affect branching patterns, nodal support, or node age estimates, whereas node ages were 1 to 4 Ma older with 5 groups than with the other grouping options, although they fell well within the respective 95% HPDs (see Figure S2).

**Figure 3.**
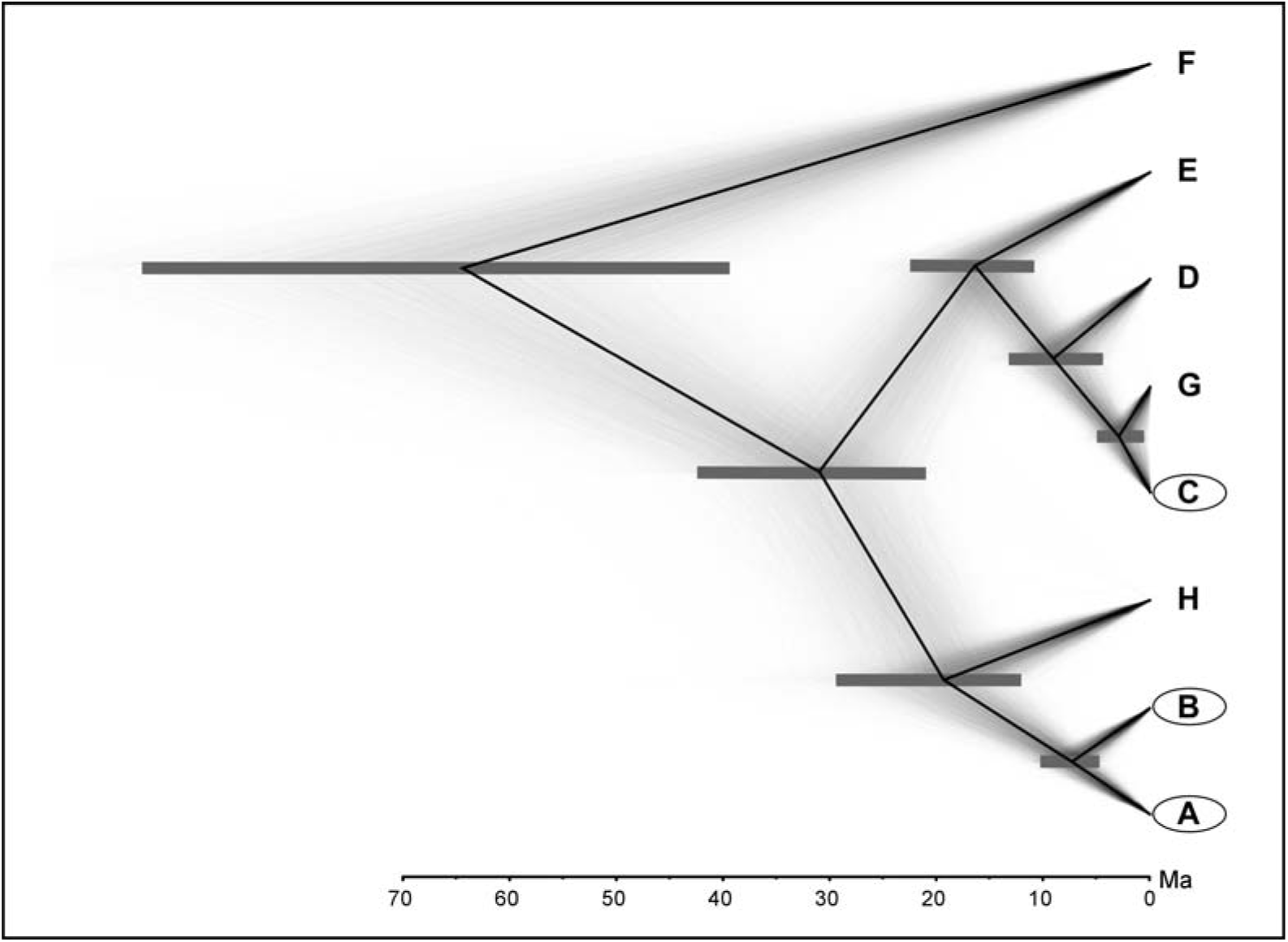
Maximum clade credibility multilocus species-tree (black line) superimposed on a cloudogram of the entire posterior distribution of species-trees obtained with *BEAST. Horizontal grey bars are 95% highest posterior densities of the estimated node ages. X-axis is in units of million years (Ma). All posterior probabilities were ≥0.99. The analysis was run with 7 terminal taxa, 5 resulting from the Bayesian analysis of the concatenated mtDNA data, plus 2 groups retrieved from the literature (clades G and H) which turned out to be the closest relatives of our insular ingroups (clades A, B, C) during previous analysis of the CO1 dataset.

## Discussion

With the ever increasing application of barcoding procedures to the study of biological diversity, a wealth of animal and plant species, including mayflies of the *Baetis rhodani* group, are revealing previously unsuspected levels and depths of variation, with potentially far-reaching implications for their taxonomy, systematics, and conservation, as well as for our understanding of their ecologies and evolutionary pathways. Unfortunately, however, in most cases mtDNA is the only component of the genetic variation analysed, which prevents a full appraisal of the biological meaning of the observed patterns of diversity (Balloux, 2010; Hudson and Coyne, 2002).

Within the *B. rhodani* group, all previous studies identified deeply divergent mtDNA (CO1) lineages, often occurring in syntopy. Nonetheless, since mtDNA provides no information about the occurrence and extent of gene exchange between diverging lineages, conclusions about the many implications of these results remained open. In the present study, we took advantage, for the first time in the *B. rhodani* group, of combined mitochondrial and nuclear perspectives, which allowed us to overcome the many pitfalls associated to the use of mtDNA alone when addressing patterns of genetic variation and species’ evolutionary histories.

Within the Tyrrhenian islands and neighbouring Elba, we found three endemic lineages (Figure S1) that were deeply divergent in their mtDNA. Indeed, the observed degree of divergence (Table 2) largely exceeded the suggested threshold for putative species, in one case (haplogroups A + B vs. C) exceeding also the one frequently observed among congeneric mayflies (18%; Ball et al., 2005). Furthermore, based on the inferred phylogenetic pattern, they appeared to be independently derived from continental relatives (see below). Nuclear DNA data confirm this pattern and, when analysed in a Bayesian population genetics framework, clearly indicate the lack of admixture between lineages, even where the lineages are sympatric (see Figure 1 and results of BAPS analysis). Thus, overall, our data clearly indicate that the three lineages we observed belong to three distinct species, under both phylogenetic and biological species concepts (Hausdorf, 2011).

Our sampling localities in northeastern Corsica (samples 18 and 19) belong to the same drainage system, and are located close to the type locality of *Baetis ingridae* (Le Bevinco, northeastern Corsica; Thomas and Soldàn, 1987). Since only lineage B was observed at these sites, it seems plausible to hypothesize that lineage B belongs to *B. ingridae*, whereas lineages A and C belong to two previously undescribed species. Nonetheless, this hypothesis deserves further confirmation in conjunction with genetic characterization of the type material.

There are several sources of uncertainty in our time estimates, including our reliance on a fixed and non-organism-specific molecular rate, and the lack of internal fossil calibration points. In addition, we refrained from using the disjunction of the plate from the continent as a dating point, since we had no data supporting this choice, and we cannot firmly exclude subsequent overseas dispersal for this mayfly (Monaghan et al., 2005). In spite of these major limitations, our dating exercise yielded a time estimate for the divergence between lineage A + B and its closest continental relative clade (19.9 Ma; see Figure 3) that fits comfortably with the oceanic spreading in the Liguro-Provençal basin (∼21–18 Ma; Gattaccecca et al., 2007; Brandano & Policicchio, 2012; Tomassetti et al., 2013), separating the Corsica-Sardinia block from the Iberian peninsula. Since intervening marine barriers may in fact fragment mayfly populations, this time coincidence appears strongly suggestive of a causal relation.

In line with previous studies (Ketmaier and Caccone, 2013), our results support an important role for the microplate disjunction and the consequent paleogeographic events in the origin of the highly endemic biota of this island complex. Nonetheless, they also emphasize that this paleogeographic process has not been the only or the major forge of endemic taxa. At least two further processes contributed: (1) speciation within the island complex, and (2) colonization from the continent.

Speciation within the island complex, which is invoked here to explain the occurrence of the divergent clades A and B, is a necessary consequence of the inferred origin of their common ancestor via microplate disjunction. Our time estimate of the divergence between clades A and B (and the associated uncertainty) would suggest a role for Corsica and Sardinia in this process. Nevertheless, in this case, we refrain from indicating this as the most plausible scenario, for three main reasons. First, there is no strong consensus on the time of final separation between Corsica and Sardinia (Alvarez, 1972; Bellon et al., 1977; Bonin et al., 1979; Gattaccecca et al., 2007). Second, both clades have been found in Corsica, and our data are not appropriate to carry out a sound analysis of the respective ancestral areas; more data will be needed for that purpose. Third, and more importantly, without clear evidence to invoke specific scenarios, we cannot exclude non-allopatric processes of speciation from the set of hypotheses (Coyne and Orr, 2004), especially considering that knowledge of the evolutionary ecology of this group of mayflies is still virtually null. Thus, how clades A and B originated, and how they reached reproductive isolation, will necessarily be a subject for future research.

Lineage C was the most differentiated among the three endemic lineages of the *B. rhodani* group on the island complex. The mean estimate of the divergence time between the respective clades (31.4 Ma) lies close to the Eocene-Oligocene transition (the so-called *Grande Coupure* or ‘great break’), an epoch of major and synchronous floral, faunal, and climatic turnover (Liu et al., 2009; Hooker et al., 2004). Nonetheless, our estimate has much uncertainty, and any hypothesis about the underling processes would require a wealth of weakly justified assumptions so that, again, we prefer not to set them at all. On the other hand, it is worth noting that both for this event and for the divergence between lineage C and its closest continental relative clade, our mean time estimates (2.9 Ma in the latter case) lie close to major and parallel climatic transitions toward cooler and drier climates. Furthermore, the Plio-Pleistocene transition (∼2.6 Ma) also introduced a marked seasonality in the Mediterranean region, and led to an abrupt sea level drop that could have made environmental conditions more favourable to a colonization of the Tyrrhenian islands from neighbouring continental areas (Thompson, 2005).

Finally, although the long polarized vicariance-dispersal debate has now been largely replaced by a more integrated view that sees them as variously interacting processes, it continues to form the basis of historical biogeographic investigations in several instances. In the context of the Tyrrhenian islands, the view that the microplate disjunction (i.e. vicariance) was the leading process responsible for the islands’ highly endemic biota has historically prevailed (Ketmaier & Caccone, 2013). Our results do not support this view, and add to some previous studies (e.g. Nascetti et al., 1996) in showing that not only both vicariance and dispersal played a part, but also that other processes, which acted within the island complex, were involved. These results thus invite us, once more (e.g. Whittaker and Fernandez-Palacios, 2007), to a sharper shift from an either-or logical framework to a more integrative one, which is increasingly appearing to provide a better perspective on the complexity of the processes that have moulded patterns of biological diversity.

## Concluding remarks

Patterns of genetic variation above and below the species level have remained largely unexplored in baetid mayflies until recent times. This study adds to several recent ones in showing that, in spite of their limited morphological variation, they usually show high levels of deeply structured genetic diversity. Besides retaining the genetic footprints of past evolutionary processes over a wide time scale, they are rather easy to sample (during the aquatic larval stages), are usually common and abundant at sites of occurrence, and given the very limited duration of the adult life stages (in the order of hours), they show comparatively limited dispersal abilities with respect to other flying insects. All these features make these organisms excellent models for historical biogeographic and evolutionary studies.

Here, we have identified the occurrence of three distinct and deeply divergent species within the *Baetis rhodani* group in the northwestern Mediterranean islands. For the time being, we refrained from making formal taxonomic descriptions, since we judged our data suboptimal for that purpose. This task shall await the examination (genetic and morphological) of the available type material, and following comparisons, the designation of new type material.

Finally, although we did not carry out statistical phylogeographic and historical demographic analyses on our samples, given the limited sample size per lineage, it did not escape our notice that the three lineages we found showed substantial intraspecific variation. Thus, as soon as a larger sample becomes available, a thorough analysis of this level of variation will allow us to shed light also on the evolutionary processes that followed settlement on the island system and the achievement of reciprocal isolation, and so will contribute to the ongoing debate concerning the relevance of intraisland microevolutionary processes.

## Acknowledgments

We are grateful to Alessandra Valentini, Simone Cardoni, Michela Paoletti, and Andrea Chiocchio, for their help with sample collection and/or with laboratory procedures. RB was supported by a post-doctoral grant from the Italian Ministry of Education, University and Research (PRIN project 2012FRHYRA). RT was supported by the INHABIT project (LIFE08 ENV/IT/000413), granted by the European Union under the LIFE + Environment Policy and Governance 2008 program.

## Data accessibility

Genbank accession numbers: KX707651-KX707786.

## Conflict of interest

The authors have no conflict of interests to declare.

## Supporting information

**Table S1.**
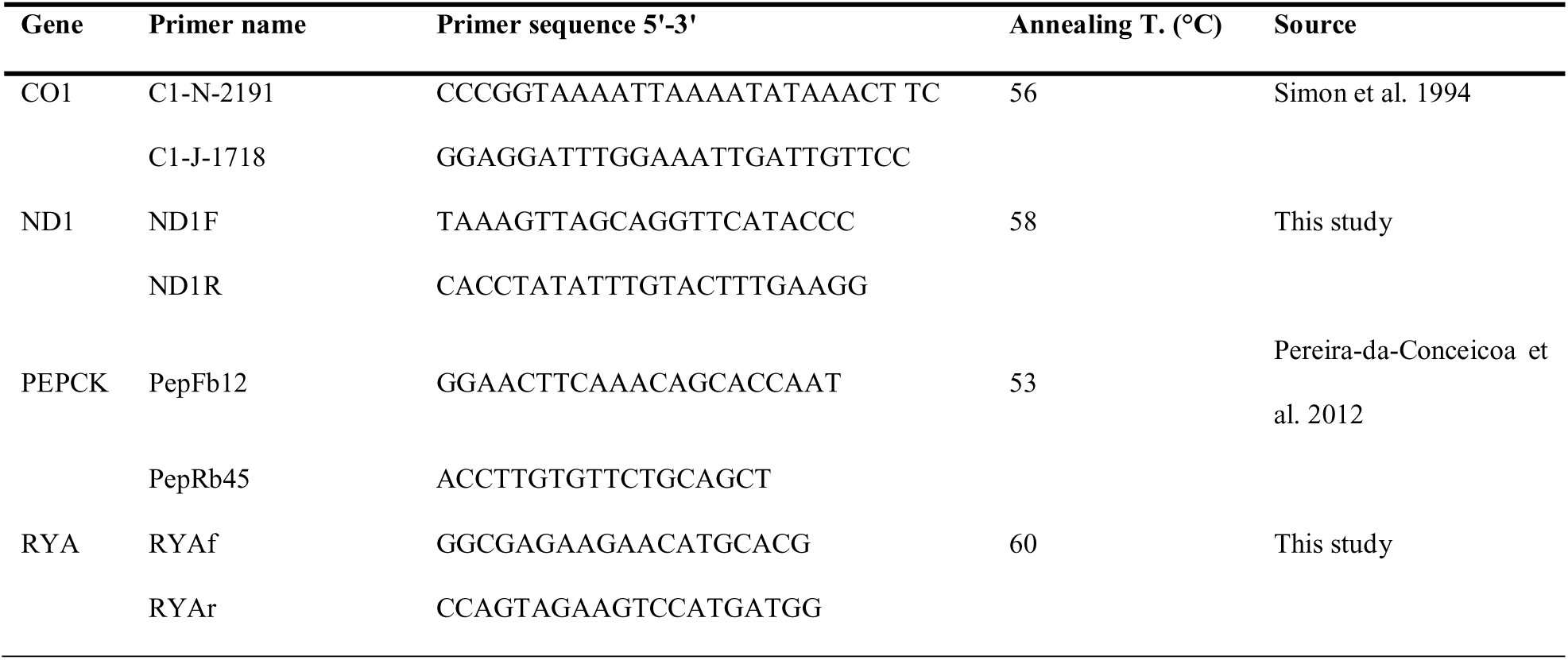
PCR primers and annealing temperatures used to amplify the two mitochondrial and the two nuclear DNA fragments used in this study.

**Figure S1.**
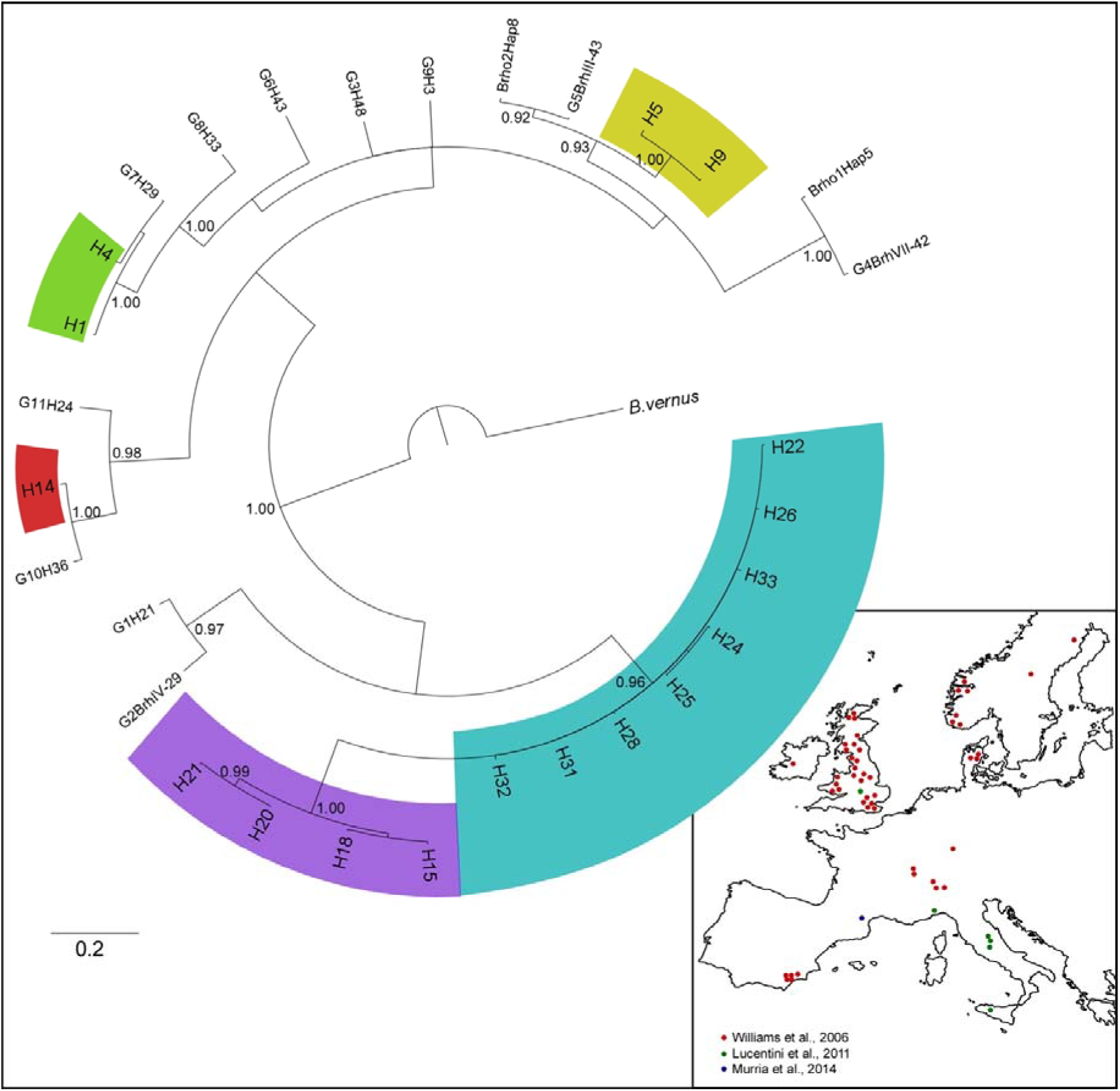
Bayesian phylogenetic tree, estimated using MRBAYES, based on the CO1 haplotypes found in this study, and 2 representatives of each main clade previously identified by Williams et al., (2006) Lucentini et al., (2011), and Murria et al., (2014). Posterior probabilities are shown at the nodes when ≥0.90.

**Figure S2.**
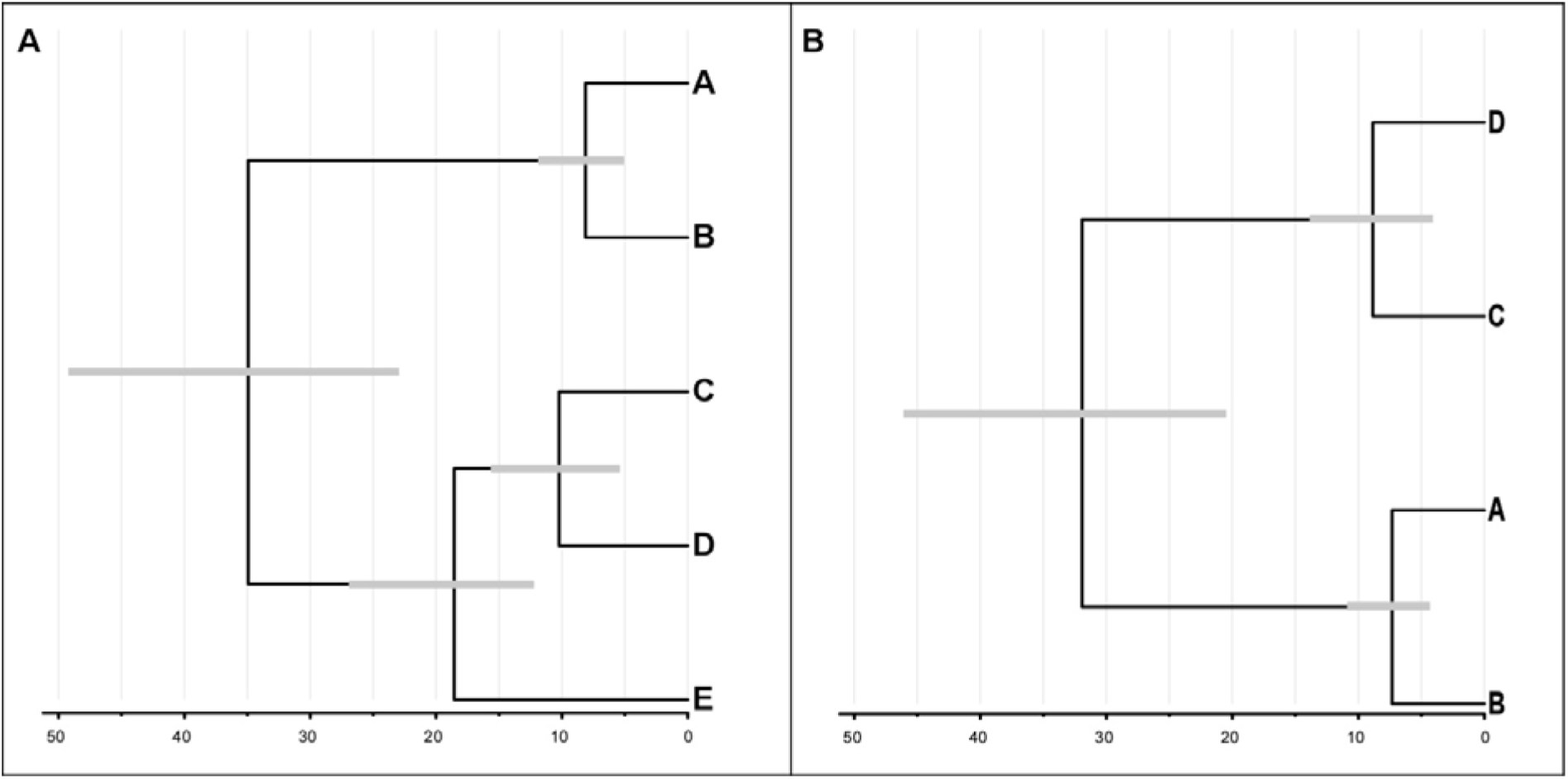
Maximum clade credibility multilocus species-trees obtained with *BEAST. Horizontal grey bars are 95% highest posterior densities of the estimated node ages. X-axis is in units of million years. All posterior probabilities were ≥0.99. (A) Analysis run with 5 terminal taxa, following results based on the concatenated mtDNA dataset (see main text Figure 1) (B) Analysis run with 4 terminal taxa, following results with BAPS based on nDNA data.

